# Effect of KNL1 on the proliferation and apoptosis of colorectal cancer cells

**DOI:** 10.1101/499889

**Authors:** Tianliang Bai, Yabin Liu, Binghui Li

**Affiliations:** Department of General Surgery, Fourth Affiliated Hospital, Hebei Medical University, Shijiazhuang, Hebei 050011, P.R. China

**Author notes:** **Corresponding author:** Professor Binghui Li, Department of General Surgery, Fourth Affiliated Hospital, Hebei Medical University, 12 Jiankang Road, Shijiazhuang, Hebei 050011, P.R. China.

**Keywords:** Kinetochore scaffold 1, colorectal cancer, cell proliferation, apoptosis

## Abstract

**Background:** To identify the expression of KNL1 in colorectal tumor tissues and to clarify the function of this gene on the proliferation capability of colorectal cancer cells.

**Methods:** Thirty-six pairs of colorectal tumor and normal tissue were collected and subjected to quantitative PCR analysis. KNL1 was under-expressed using lentiviral transfection, and the function of KNL1 was evaluated by proliferation assay, colony formation and apoptosis assay.

**Results:** KNL1 was highly expressed in colorectal tumor tissues. KNL1 downregulation inhibited colorectal cancer cell proliferation and promoted apoptosis.

**Conclusions:** KNL1 plays an effective role in decreasing the apoptosis and promoting the proliferation of colorectal cancer cells.

## Introduction

Colorectal carcinoma (CRC) has been reported to be the second deadliest cancer ^1-4^ Although the overall morbidity and mortality of CRC has decreased in recent decades, there has been a considerable increase in patients <50 years of age ^4-6^. Additionally, the 5-year survival of CRC is only 14% as the tumor is often not completely resectable^5, 6^. Despite advances in therapeutic strategies and diagnostic approaches, the prognoses for patients with CRC typically remain poor ^7^. Therefore, there is an urgent need to develop new CRC treatment approaches. We previously reported the effects of D40 cloning on chromosome 15 ^8^, and D40 was characterized to be a cancer-related gene that is primarily expressed in the testis under normal conditions, but is widely expressed in primary tumors of various origins as well as assorted cancer cell lines ^9, 10^. A previous study has reported the mutual translocation of AF15q14 with the MLL (Mixed lineage leukemia) gene in acute myeloid leukemia ^11^. AF15q14, a human gene on chromosome 15, was also identical to the D40 gene ^8^. Furthermore, D40 expression has been reported to be associated with the pathological features of primary lung tumors, such as degree of differentiation, and patients’ smoking history ^10^. In a study of the testes, D40 was found to be expressed in pre-acrosomes and spermatocytes ^12^. A follow up study revealed that blinkin, a type of kinetochore protein in the mitotic machinery was identical to KNL1. In addition to binding of KNL1/blinkin with Ndc80 and Mis12 complexes, of which the KMN network is comxsprised, binding is also observed between KNL1/blinkin and other proteins, including tubulin, spindle assembly checkpoint (SAC) proteins BubR1 and Bub1, and Protein Phosphatase 1 ^13-17^. Such binding suggests that KNL1/blinkin has crucial impacts on kinetochore formation in terms of the connection between spindles and chromosomes, as well as the regulation of SAC ^13-18^.

Primary microcephaly (MCPH) is a rare congenital neurodevelopment disorder that is known to have autosomal recessive features^19, 20^. Due to different levels of mental retardation, the head circumference of MCPH patients will be reduced and their brains are smaller. However, additional somatic or neurological disorders are not generally observed in patients with MCPH. MCPH is genetically heterogeneous and is controlled by several chromosomal loci are responsible ^19^. In previous studies, an MCPH locus has been reported on chromosome 15 ^20^, and the gene responsible has been identified as CASC5, which is also identical to KNL1 ^21^. Considering that brain development requires the rapid division of cells, this suggests that KNL1 is significant for *in vivo* cell division.

In the present study, we analyzed KNL1 expression in various CRC and matched adjacent noncancerous tissues, as well as in CRC cell lines. We also studied how KNL1 knockdown impacts CRC cells *in growing under an in-vitro in vitro* by transfecting the RKO and HT-29 CRC cell lines with shKNL1 or shCtrl.

As described previously ^8, 10, 11, 13, 16^, KNL1 has a number of alternative names, including AF15q14, KIAA1570, CASC5, blinkin and D40. The nomenclature KNL1 was used in the present study.

## Materials and methods

### Patients and samples

We collected human CRC and matching adjacent noncancerous tissue samples from 36 patients at the Department of General Surgery of the Fourth Hospital of Hebei Medical University (Shijiazhuang, Hebei, China). All patients consented to inclusion in the present study and all experiments were approved by the Ethics Committee of the Fourth Hospital of Hebei Medical University.

### Cell lines and cell culture

We purchased the human CRC cell lines (SW480, HT-29, HCT-116 and RKO) and the normal colonic epithelial cell line NCM460 from the American Type Culture Collection (ATCC; Manassas, VA, USA). Cells were maintained in RPMI-1640 (Gibco; Thermo Fisher Scientific, Inc., Waltham, MA, USA) supplemented with 10% fetal bovine serum (Gibco; Thermo Fisher Scientific, Inc.) in a humidified atmosphere containing 5% CO_2_ at 37°C.

### RNA extraction and reverse transcription-quantitative polymerase chain reaction (RT-qPCR)

We used TRIzol^®^ reagent (Invitrogen; Thermo Fisher Scientific, Inc.) to extract whole RNA from the sampled cells and tissues. RT was performed to generate cDNA using M-MLV Reverse Transcriptase (Promega Corp., Madison, WI, USA) according to the manufacturer’s protocol. A total of 1 μl cDNA was used as the template for qPCR. GAPDH was used as an internal control. Primer sequences were as follows: KNL1, forward 5′-ACA TTG GAA AAA GCG CAA GTTG-3′ and reverse 5′-TTG CAC TGG GCA ATA ATT GGC-3′; GAPDH, forward 5′-TGA CTT CAA CAG CGA CAC CCA-3′ and reverse 5′-CAC CCT GTT GCT GTA GCC AAA-3′. Thermocycling comprised initial denaturation at 95°C for 30 sec followed by 45 cycles of denaturation at 95°C for 5 sec and extension at 60°C for 30 sec. The PCR products for GAPDH and KNL1 were 121 and 213 bp in length, respectively. Each experiment was performed in triplicate and relative quantitative gene expression was calculated as previously described ^22^.

### Western blotting

Cells and tissues were lysed using RIPA buffer (Thermo Fisher Scientific, Inc.). Lysates were electrophoresed by 10% SDS-PAGE (Thermo Fisher Scientific, Inc.) and transferred onto PVDF membranes (Thermo Fisher Scientific, Inc.). After blocking for 1 h in 5% skimmed milk, membranes were incubated overnight at 4°C with rabbit antibodies against KNL1 (dilution, 1:500; Biorbyt Ltd., Cambridge, UK) and GAPDH (dilution, 1:800; Santa Cruz Biotechnology, Inc., Dallas, TX, USA). TBST was used to wash all the membranes 4 times. Membranes were subsequently incubated at room temperature for 1 h with an HRP-conjugated secondary antibody (dilution, 1:1,0000; catalog no., 323-065-021; Jackson ImmunoResearch, Inc., West Grove, PA, USA). Protein bands were visualized using a chemiluminescence system (Thermo Fisher Scientific, Inc.) according to the manufacturer’s protocol. The band intensities were quantified using Image-Quant Software v3.0 (LI-COR Biosciences). This experiment was repeated in triplicate.

### Construction of recombinant lentiviral vector and cell transduction

KNL1-shRNA and control-shRNA were designed by GeneChem Co., Ltd. (Shanghai, China) from the unabridged sequence of D40/KNL1, which is targeted at the human KNL1 gene (Genbank no. NM_144508). The targeting sequence of KNL1 was 5’-CAG AGT TGT ATG GTG GAAA-3’. To test how efficient KNL1 knockdown was, stem-loop oligonucleotides were synthesized and inserted into a lentivirus-based pGV115-GFP vector (GeneChem Co. Ltd.) with AgeI/EcoRI sites. Lentivirus particles were prepared as previously described ^23^. RKO and HT-29 cells (2×10^5^ cells/well) were cultured in 6-well plates and transfected with either negative control (shCtrl) lentivirus or KNL1 (shKNL1) lentivirus at a multiplicity of infection (MOI) of 5. Cells were incubated for 5 days at 37°C in an atmosphere containing 5% CO_2_. After 72 h of transfection, cells were observed using a fluorescence microscope (IX71; Olympus Corp., Tokyo, Japan). When the 5-day transfection was complete, knockdown efficiency was measured using western blotting and RT-qPCR.

### Cell growth assay

ShKNL1-transfected or shCtrl-transfected RKO and HT-29 cells in the logarithmic growth phase were cultured in 96-well plates at a density of 2,000 cells per well for 5 days at 37°C in an atmosphere containing 5% CO_2_. Cell numbers were recorded every day using a Celigo Imaging Cytometer (Nexcelom Bioscience, Lawrence, MA, USA), and all were performed in triplicate.

### Methyl-thiazol-tetrazolium (MTT) assay

ShKNL1-transfected or shCtrl-transfected RKO and HT-29 cells were again seeded and cultured in 96-well plates at a density of 2000 cells/well at 37°C. Cell proliferation was assessed each day for 5 days. Briefly, 20 μl MTT (5 mg/ml, Genview, Craigie Burn, Australia) was added to each well and incubated at 37°C for 4 h. The culture medium was removed and 100 μl of dimethyl sulfoxide was added to each well to dissolve the crystals. Plates were agitated for 2-5 min and the absorbance at 490 nm was read using a microplate reader (Tecan infinite, Tecan GmbH, Austria). All experiments were performed in triplicate.

### Colony formation assay

Cells transfected with shKNL1 or shCtrl vectors in the logarithmic growth phase were re-suspended in RPMI-1640 and seeded in 6-well plates at a density of 800 cells/well. Cells were incubated for 14 days and fluorescence microscopy (IX71; Olympus Corp.) was used to observe cell colonies. Cells were fixed with paraformaldehyde (1 ml/well; Sangon Biotech Co. Ltd., Shanghai, China) for 30 min. Cells were subsequently washed with PBS and stained with 500 μl 0.5% crystal violet (Sangon Biotech Co. Ltd., Shanghai, China) for 5 min. After the cellular staining, ddH_2_O was used to wash the stained cells, which were then dried at room temperature. Finally, cell colonies were observed under a microscope (Cai Kang Optical Instrument Co., Ltd, Shanghai, China). All experiments were performed in triplicate.

### FACS analysis. Fluorescence-activated cell sorting (FACS) was used to analyze cell apoptosis^24^

In brief, RKO cells transfected with shCtrl or shKNL1 were seeded in 6-well plates for 120 h and grown to 70% confluence, harvested and washed with 1X binding buffer, following which 10 μl of Annexin V-allophycocyanin (APC) Detection kit (eBioscience; Thermo Fisher Scientifc, Inc.) was added per 200 μl of the adjusted suspension for 15 min in the dark at room temperature. Flow cytometry analysis after cellular staining. All experiments were performed in triplicate.

### Statistical analysis

SPSS version 19.0 for Windows (IBM Corp., Armonk, NY, USA) was used for statistical analysis and all data are presented as the mean ± standard deviation. Comparisons between the two groups were made using Student’s t-test and P<0.05 was considered to indicate a statistically significant difference.

## Results

### Compared with normal adjacent tissues, the expression of KNL1 was obviously higher in colorectal tumor tissues

RT-qPCR and Western blot were used to measure the expression levels of KNL1 in CRC and para-cancerous tissues. The results revealed that, compared with normal colorectal tissues, KNL1 mRNA and protein levels were much higher in CRC tissues (Fig. 1A and B).

**Figure 1.**
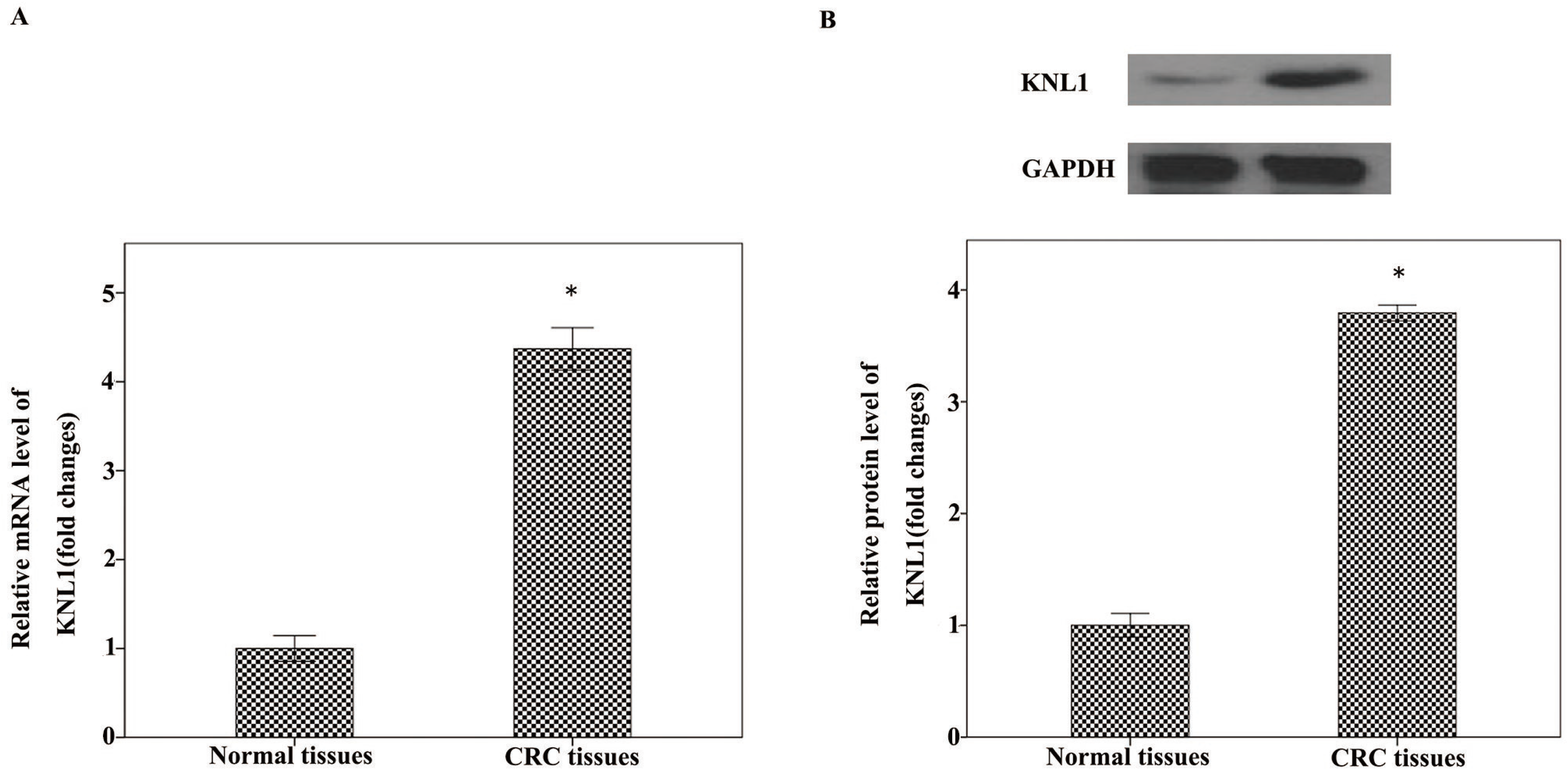
Expression of KNL1 is upregulated in CRC tissues. The mRNA (A) and protein (B) expression levels of KNL1 were significantly higher in CRC tissues than in the corresponding noncancerous tissues.*p<0.05.

### KNL1 expression in 4 colorectal cancer cell lines

RT-qPCR and western blotting were used to measure the expression of KNL1 mRNA and protein in SW480, RKO, HT-29, HCT-116 CRC cells and the normal colonic epithelial cell line NCM460. The results indicated that KNL1 mRNA and protein was highly expressed in in SW480, RKO, HT-29 and HCT-116 cell lines (Fig. 2).

**Figure 2.**
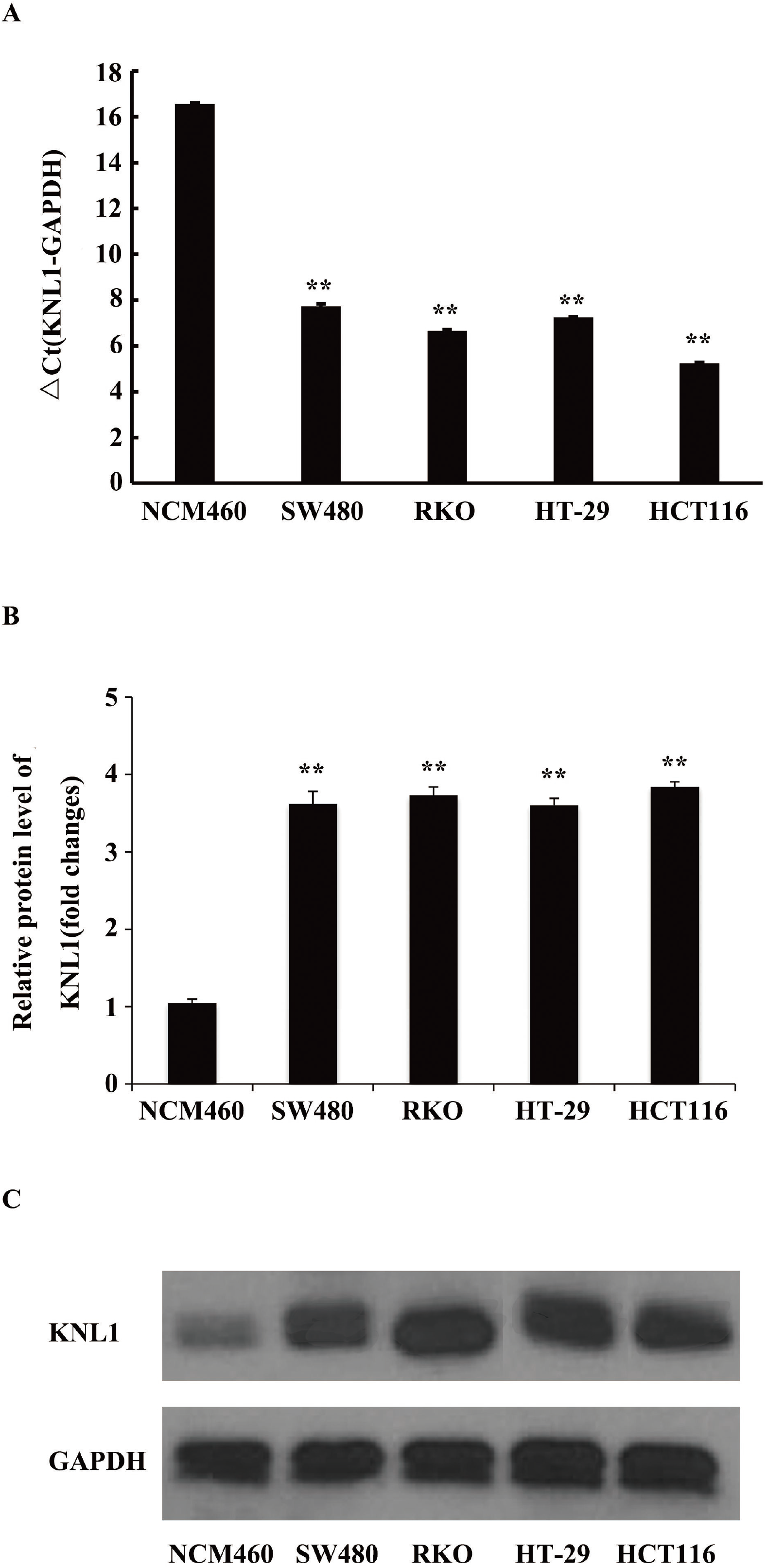
KNL1 mRNA and protein levels in four colorectal cell lines and the normal colonic epithelial cell line. (A) KNL1 mRNA expression levels determined by RT-qPCR. (B and C) KNL1 protein expression levels determined by western blotting. **P <0.01.

### Lentivirus-mediated knockdown of KNL1 in CRC cells

To assess the mechanism of KNL1 in CRC, KNL1 expression was knocked down in RKO and HT-29 cells using transfection. After 3 days, >80% of the cells were successfully transfected with either an shCtrl or an shKNL1 lentivirus (Fig. 3A and D). RT-qPCR was performed 5 days after transfection and KNL1 mRNA expression in the shKNL1 group was much lower compared with in shCtrl-infected control cells (Fig. 3B and E). These results were confirmed using western blotting, and it was clear that shKNL1-transfected cells had largely reduced KNL1 levels, indicating that the target gene was successfully knocked down (Fig. 3C and F).

**Figure 3.**
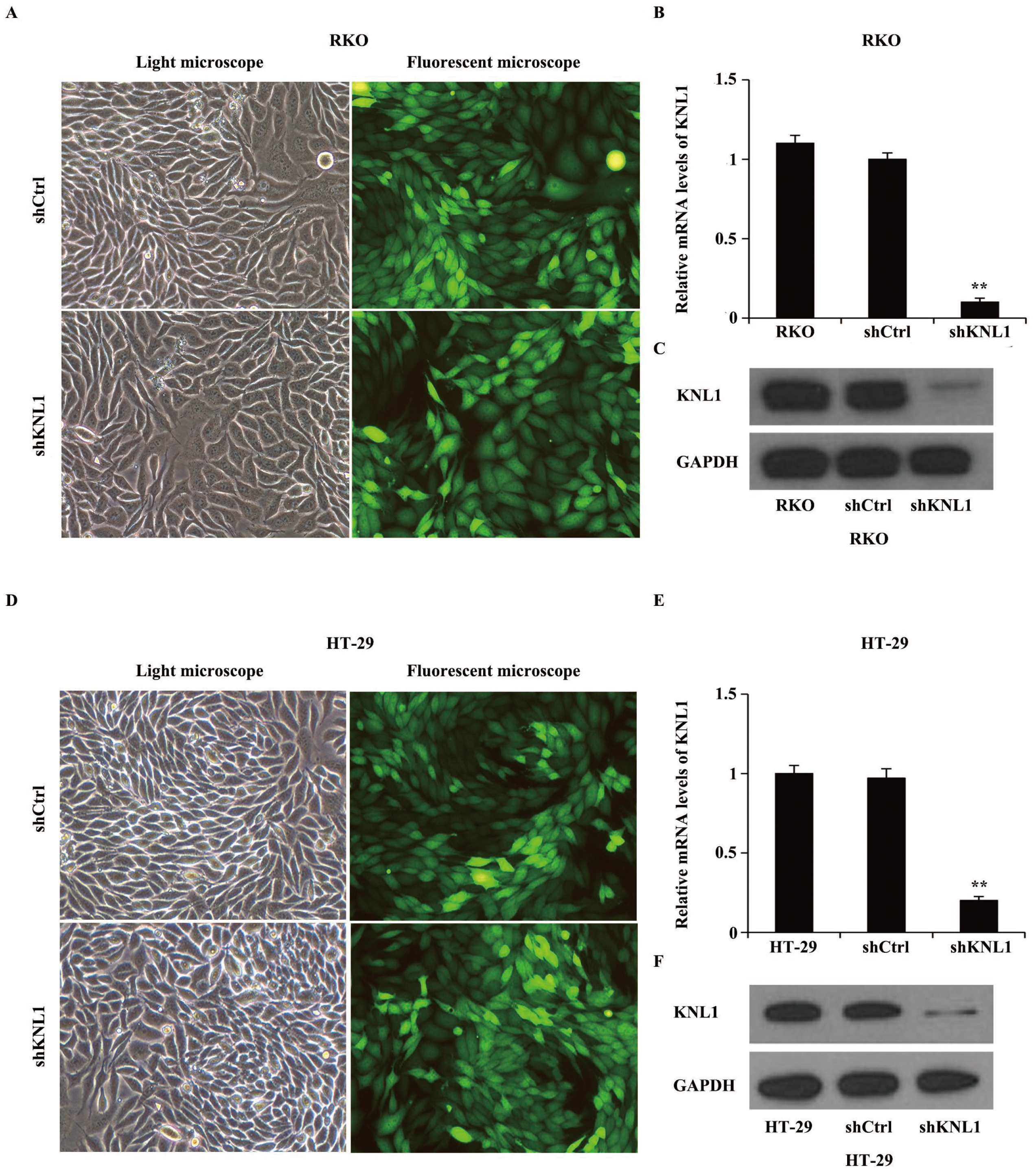
Knockdown of KNL1 in RKO and HT-29 cells infected with shKNL1 or shCtrl. (A and D) Cells were examined by fluorescent and light microscopy at day 3 post-infection (x200, magnification). Representative images of the cultures are shown. (B and E) KNL1 mRNA levels were analyzed using RT-qPCR at day 5 post-infection. KNL1 mRNA level decreased significantly after KNL1 knockdown (**P<0.01). (C and F) KNL1 protein expression was analyzed by western blotting in the transduced RKO and HT-29 cells.

### The CRC cell proliferation is inhibited by KNL1 knockdown

RKO and HT-29 cells transfected with shCtrl or shKNL1 lentivirus were seeded into 96-well plates and analyzed using Celigo for 5 consecutive days. The number of cells in the shCtrl group considerably increased over the 5 days, whereas the cell number in the shKNL1-transduced group increased slightly (Fig. 4A-C and 4F-H). These results suggest that KNL1 knockdown significantly inhibits CRC cell proliferation, which was further confirmed by the MTT results (Fig. 4D-E and 4I-J). It appears that KNL1 expression determines the growth of RKO and HT-29 cells.

**Figure 4.**
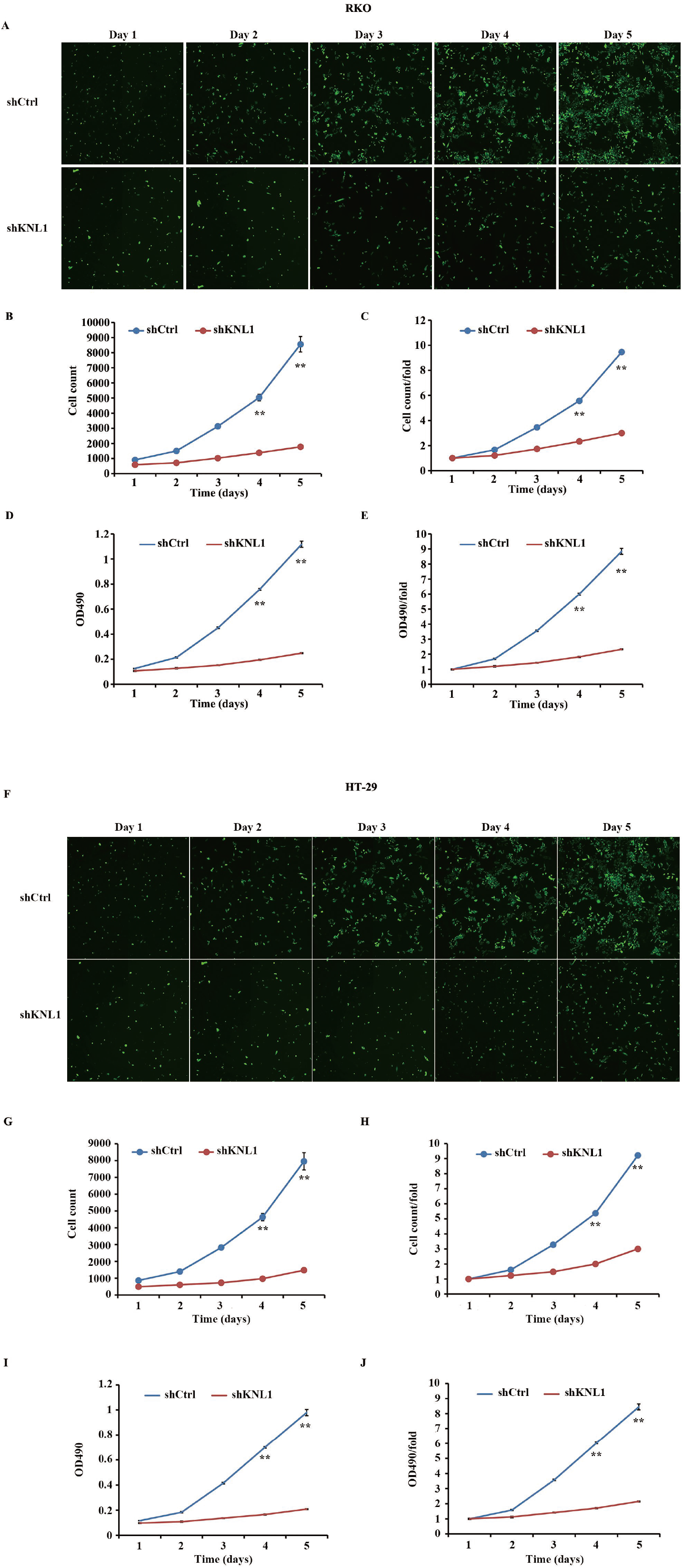
KNL1 knockdown inhibits RKO cell growth as compared with control group. (A) RKO cells are infected by shCtrl and shKNL1 for 5 days, and fluorescence microscope (green staining) shows that the cell count is increased in a time-dependent model. (B) Celigo Cell Counting indicated that cell growth is slower in shKNL1 group than in shCtrl group. (C) The cell count and count fold curves show significant differences between shKNL1 group and shCtrl group. (D) The proliferative abilities of RKO cells transfected with shCtrl and shKNL1, which were evaluated by MTT assay. (E) Downregulation of KNL1 remarkably inhibited the proliferation rate of RKO cells as measured by the MTT assay. KNL1 knockdown inhibits HT-29 cell growth as compared with control group. (F) HT-29 cells are infected by shCtrl and shKNL1 for 5 days, and fluorescence microscope (green staining) shows that the cell count is increased in a time-dependent model. (G) Celigo Cell Counting indicated that cell growth is slower in shKNL1 group than in shCtrl group. (H) The cell count and count fold curves show significant differences between shKNL1 group and shCtrl group. (I) The proliferative abilities of HT-29 cells transfected with shCtrl and shKNL1, which were evaluated by MTT assay.(J) Downregulation of KNL1 remarkably inhibited the proliferation rate of HT-29 cells as measured by the MTT assay. **P <0.01.

### The formation of CRC cell colonies is inhibited by the KNL1 knockdown

RKO and HT-29 cells were stained with crystal violet to assess colony formation (Fig. 5A and C). The number of the cells per colony was significantly higher in the shCtrl group compared with the shKNL1 group (shCtrl: 229±9 vs. shKNL1: 40±5; shCtrl: 259±2 vs. shKNL1: 55±3; P<0.01) (Fig. 5B and D), indicating that the endogenous reduction in KNL1 expression significantly inhibited CRC cell growth.

**Figure 5.**
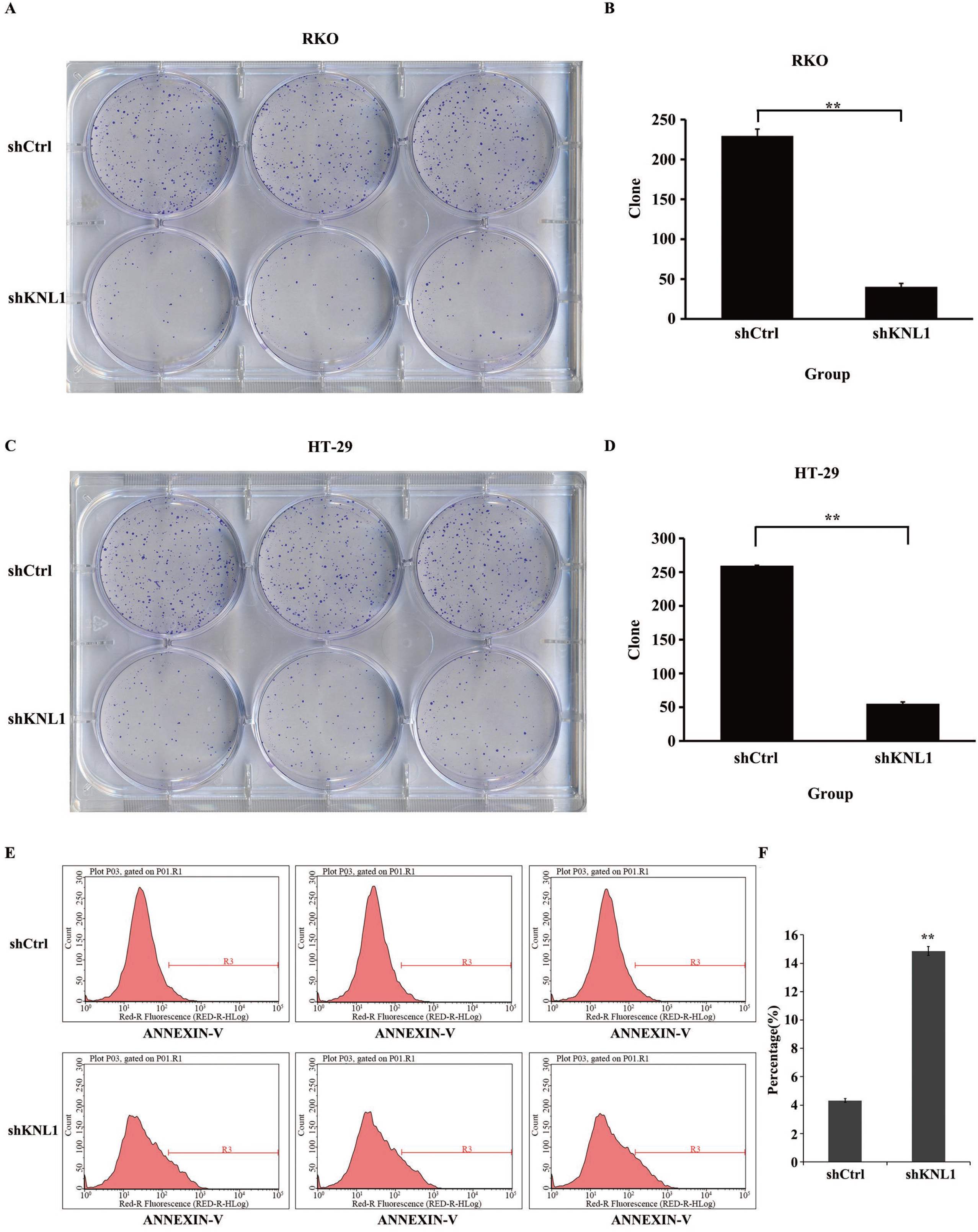
KNL1 silencing represses RKO and HT-29 cells colony formation. (A and C) Photomicrographs of crystal violet-stained colonies of RKO and HT-29 cells growing in 6-well plates for 10 days after infection. (B and D) The number of cells in each colony of RKO and HT-29 cells was counted. Cell number in shKNL1 group was significantly fewer relative to shCtrl-transduced cells. RKO cell apoptosis analyzed by flow cytometry. (E) Cells are transfected with shKNL1 and shCtrl, apoptosis is analyzed by flow cytometry. Each group is shown in triplicates. (F) The resulting data indicate that the cell apoptosis is significantly higher in shKNL1 group than in shCtrl group. **P <0.01.

### KNL1 knockdown affects the apoptosis of RKO cells

The apoptotic rate of RKO cells in the shKNL1 group was 14.86±0.30%, whereas it was 4.32±0.13% in the shCtrl group. The results suggest that apoptosis was significantly increased in the shCtrl group compared with the shNKL1 group (P<0.05) (Fig. 5E and F).

## Discussion

The morbidity and mortality rate of CRC has risen rapidly, and it has attracted worldwide attention^25, 26^. Despite improvements in clinical practice and treatment, outcomes to CRC patients are still poor due to metastasis or drug resistance^27, 28^. Therefore, it is necessary to determine new therapeutic targets and to better understand the underlying molecular mechanism.

The occurrence and development of tumors are associated with both behavioral and environmental factors, and tumorigenesis results from genetic mutations accumulating over time. Identifying the affected genes may allow for the development of advanced diagnostic and treatment methods for cancer.

The spindle checkpoint controls mitotic progression. The human kinetochore oncoprotein AF15q14/KNL1, which belongs to the Spc105/Spc7/KNL-1 family, has been reported to provide direct links between Bub1 and BubR1 spindle checkpoint proteins and the kinetochores, and also plays a crucial role in the alignment of chromosomes and spindle checkpoints. KNL1 RNAi gives induces accelerated mitosis due to chromosome misalignment and checkpoint failure, which are caused by the lack of microtubule attachment and kinetochores^16, 29^. Another study reported that transfection with KNL1 siRNA resulted in cancer cell apoptosis and inhibited the growth of these cells in both *in vitro* and *in vivo*, regardless of p53 status was ^30^. However, to the best of our knowledge, there have not previously been specific studies regarding KNL1 expression in human cancers, particularly CRC.

The results of the present study suggest an obvious relationship between KNL1 and CRC cell apoptosis and proliferation. Although many studies described the abnormal mitosis in cancer cells treated with KNL1 siRNA^16, 31^, we are the first to demonstrate that KNL1 downregulation induced apoptosis and inhibited the growth of CRC cell lines. However, there were a number of limitations to consider. The number of clinical samples was small and a larger sample size is required to verify the expression of KNL1 in colorectal tumors. There was also a lack of relevant clinical information regarding the study participants.

In conclusion, the results of the present study suggest that shRNA downregulates the expression of KNL1 in CRC cells, which in turn induces apoptosis and inhibits proliferation. KNL1 knockdown may therefore be an effective assumed therapeutic approach for the treatment of CRC in which KNL1 is overexpressed.

## Acknowledgement

Not applicable.

## Conflict of interest

All authors declare that they have no conflict of interest.

## Availability of data and materials

The datasets used and/or analyzed during the current study are available from the corresponding author on reasonable request.

